# YOTO (You Only Think Once): A Human EEG Dataset for Multisensory Perception and Mental Imagery

**DOI:** 10.1101/2025.04.17.645384

**Authors:** Yan-Han Chang, Hsi-An Chen, Min-Jiun Tsai, Chun-Lung Tseng, Ching-Huei Lo, Kuan-Chih Huang, Chun-Shu Wei

**Affiliations:** Department of Computer Science, National Yang Ming Chiao Tung University, Hsinchu, Taiwan; Interdisciplinary Science Degree Program, National Yang Ming Chiao Tung University, Hsinchu, Taiwan; Institute of Mathematical Modeling and Scientific Computing, National Yang Ming Chiao Tung University, Hsinchu, Taiwan; Institute of Education, National Yang Ming Chiao Tung University, Hsinchu, Taiwan; Brain Science and Technology Center, National Yang Ming Chiao Tung University, Hsinchu, Taiwan; Institute of Biomedical Engineering, National Yang Ming Chiao Tung University, Hsinchu, Taiwan

## Abstract

The YOTO (You Only Think Once) dataset presents a human electroencephalography (EEG) resource for exploring multisensory perception and mental imagery. The study enrolled 26 participants who performed tasks involving both unimodal and multimodal stimuli. Researchers collected high-resolution EEG signals at a 1000 Hz sampling rate to capture high-temporal-resolution neural activity related to internal mental representations. The protocol incorporated visual, auditory, and combined cues to investigate the integration of multiple sensory modalities, and participants provided self-reported vividness ratings that indicate subjective perceptual strength. Technical validation involved event-related potentials (ERPs) and power spectral density (PSD) analyses, which demonstrated the reliability of the data and confirmed distinct neural responses across stimuli. This dataset aims to foster studies on neural decoding, perception, and cognitive modeling, and it is publicly accessible for researchers who seek to advance multimodal mental imagery research and related applications.

## Introduction

Mental imagery is an important component of human cognition and has value in many application domains. Mental imagery refers to an internal process in which perception-like representations of objects, scenes, events, or sensations emerge without direct external inputs ^1^. Studies suggest that these internally generated representations activate neural substrates that also respond to actual perception, indicating that mental imagery functions as a neural simulation of real sensory experiences ^2,3^.

Among the various types of mental imagery, visual imagery has been studied in a more extensive way ^4^. Previous research has shown that visual imagery activates early visual cortical areas (V1–V4) and is closely associated with spatial reasoning, visual memory, and visual creativity ^5,6^. Auditory imagery engages the auditory cortex (A1) ^7^, which plays an essential role in music memory, language comprehension, and speech learning processes. Motor imagery, associated with the motor cortex (M1), is frequently used in athletic training ^8^, motor skill enhancement, and rehabilitation therapies to improve muscle coordination and precision of movement ^9^. Beyond these widely studied types, olfactory imagery involves the olfactory cortex and related limbic structures, such as the piriform cortex and amygdala, and can trigger emotional and autobiographical memory activation ^10,11^. Tactile imagery helps mentally reconstruct the sense of touch, including temperature, texture, and pressure ^12^. Gustatory imagery activates the insular cortex and is closely related to emotional regulation and appetite control ^13^. Emotional imagery involves emotional processing regions, including the amygdala and anterior cingulate cortex (ACC), and has been applied effectively in treatments for psychological disorders such as post-traumatic stress disorder (PTSD) and anxiety disorders ^14,15^. Importantly, different types of mental imagery often co-occur in naturalistic settings. Neuroimaging findings suggest that these internally generated sensations partially recruit the same neural substrates that process real sensory input ^1,16^. Mental imagery, in essence, is not merely symbolic or abstract representation; rather, on multiple levels, it simulates the brain’s response to actual perception. However, mental imagery does not occur exclusively in task-specific states; even in the absence of external stimuli, the brain remains highly active during resting states, particularly in the medial prefrontal cortex, the posterior cingulate cortex / precuneus, and the lateral cortical regions, which consistently exhibit coordinated activity during rest ^17,18^. This network, known as the default mode network (DMN), is closely related to memory recall ^19^, mind wandering and daydreaming ^20^, memory retrieval, self-referential thought, and mental simulation ^21–23^.

Researchers have applied neural representations of mental imagery in electroencephalography (EEG) decoding technologies. The neural mechanisms of mental imagery include sensory simulation, memory retrieval, and internal thought regulation, and these mechanisms illuminate how the brain reconstructs sensory experiences in the absence of external inputs ^4,24^. Studies indicate that mental imagery produces measurable EEG patterns, such as alpha-wave changes and altered occipital gamma waves ^25,26^. These neural markers suggest that EEG signals offer a promising means to recognize and classify mental imagery without relying on external stimuli. Visual imagery appears in alpha-wave modulation and changes in gamma-band activity in occipital regions ^25,26^. Auditory imagery involves cortical areas linked to auditory processing. Motor imagery modulates sensorimotor rhythms and aids rehabilitation and motor control. Each imagery type reflects distinct neural patterns, and EEG signals detect these patterns in real time. The ability to capture such activity advances interactive technologies that integrate mental imagery for communication or control. Most studies focus on unimodal sensory imagery, such as purely visual or auditory forms. Research on multisensory integration remains less extensive ^27^. Concurrent processing of visual, auditory, and tactile imagery needs better understanding of neural mechanisms that integrate these modalities. Mental imagery EEG datasets also encounter technical constraints. A low signal-to-noise ratio (SNR) and high interindividual variability limit reproducibility of research findings ^28^. Methods lack uniform standards, which complicates generalization of decoding results ^29^. Observing EEG signals during mixed imagery tasks may clarify how the brain coordinates and merges several internal simulations.

The YOTO (You Only Think Once) dataset holds strong potential for advancing research into the neural mechanisms of mental imagery and resting-state brain activity. It provides a rich collection of non-invasive EEG recordings during multimodal mental imagery tasks and spontaneous rest from a diverse group of participants. We anticipate a wide range of applications for this dataset. For instance, it can be used to develop and evaluate EEG-based decoding models of imagined sensory experiences, investigate and compare dynamics of perceived and imagined sensory responses, and explore the integration of multisensory representations. In sum, YOTO complements existing EEG resources by providing a high-quality, systematically curated dataset tailored to the study of internal mental states and multimodal cognitive process.

## Methodology

### Participants

26 healthy participants (16 males, 10 females) volunteered to participate in the study, with a mean age of 23.3 years (median: 23 years, range: 20–36 years). All participants had normal or corrected-to-normal vision and provided their written informed consent prior to the experiment. Exclusion criteria included screen-induced dizziness, major diseases, irregular sleep patterns, poor sleep quality, disability, psychiatric disorders, or pregnancy. This study was approved by the Human Subjects Research Ethics Committee of the National Chiao Tung University (Approval No. NCTU-REC-108-128F).

### EEG Data Acquisition

EEG signals were recorded in an electromagnetically shielded chamber using a high-fidelity electrophysiological recording system. The Polhemus 3SPACE FASTRAK system was used to position the Cz reference point before securing the EEG headset (Figure 1). Thirty-two electrodes, including two reference electrodes at A1 and A2, were placed according to the 10-20 international system. Thirty-channel EEG signals were amplified and transmitted to a computer via a SynAmps RT 64 channel amplifier (Compumedics Neuroscan), digitized at 1000 Hz, and event markers were transmitted via a parallel port to indicate the onset of the trial, stimulus presentation, resting phase, and imagery phases.

**Fig. 1:**
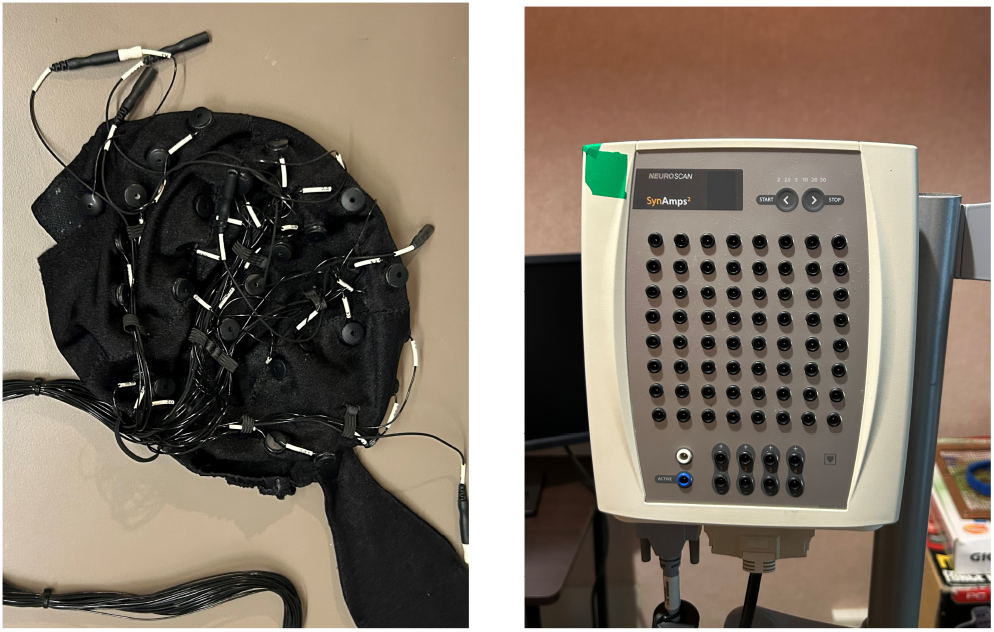
EEG Data Acquisition System Architecture. (a) The EEG cap with a 30-channel electrode configuration, (b) The 64-channel high-resolution EEG signal amplifier.

### Experimental Protocol

Each participant completed two separate sessions on different days, each session consisting of four blocks of 48 trials, interspersed with short breaks(Figure 2). During the experiment, participants were instructed to keep fixation on a central cross while participating in mental imagery tasks associated with the stimuli presented.

**Fig. 2:**
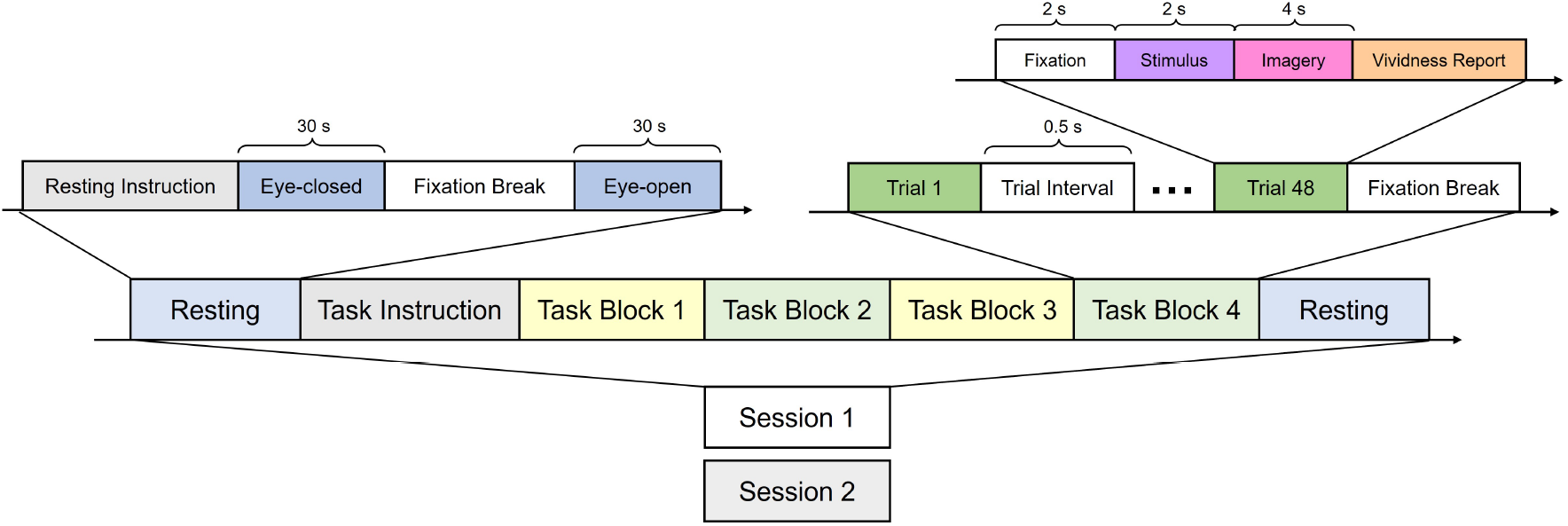
Experimental Procedure and Trial Structure. Detailed presentation of the trial structure, clearly illustrating task phases including fixation, stimulus presentation, imagery phase, and subjective self-report.

Each trial followed a structured sequence, as shown in Table 1, beginning with a fixation period (2 s), during which the participants focused on a central cross to clear their thoughts. This was followed by a stimulus presentation phase (2 s), in which a visual, auditory, or combined stimulus was randomly displayed. Subsequently, the participants entered the imagery phase (4 s), during which they mentally visualized the stimulus received previously. Finally, during the self-report phase, participants rated the vividness of their mental imagery on a scale from 1 to 5. The duration of this phase varied across individuals. To assess baseline neural activity, resting-state EEG data were recorded both before and after the experimental session.

**Table 1:** Experimental procedure of a single trial consisting of four sequential phases.

| Phase | Duration | Description |
| --- | --- | --- |
| Fixation | 2 s | Participants fixated on a central cross to prepare for the task. |
| Stimulus Presentation | 2 s | A randomly selected visual, auditory, or multimodal stimulus was presented. |
| Imagery Phase | 4 s | Participants imagined the stimulus they had just perceived. |
| Self-Report | Varies | Participants rated the vividness of their mental imagery. |

### Stimuli Details

In the imagery task, visual and auditory stimuli were presented, individually or in combination, to ensure a complete examination of sensory processing.

The visual stimuli, as illustrated in Figure 3, consisted of a gray square and two facial images (one male, one female) ^30^, while the auditory stimuli included three human short vowels (/a/, /i/, /o/) and three piano tones (C: 261.63 Hz, D: 293.66 Hz, E: 329.63 Hz). These stimuli were systematically combined to create 27 unique stimulus conditions, which encompassed three categories: visual-only, auditory-only, and multimodal (visual-auditory) stimuli. To ensure a balanced distribution of stimuli in trials, a weighted randomization strategy was implemented (Table 2, Trials/Block). This weighted randomization approach prevented overrepresentation of specific conditions while maintaining sufficient exposure to all stimulus types, ensuring a balanced and unbiased experimental design.

**Table 2:** Stimuli types and their corresponding trials per block.

| Modality | Stimuli | Trials / Block |
| --- | --- | --- |
| **Visual** | Square | 6 |
|  | Male face | 3 |
|  | Female face | 3 |
| **Auditory** | Vowel /a/ | 2 |
|  | Vowel /o/ | 2 |
|  | Vowel /i/ | 2 |
|  | Tone C (262 Hz) | 2 |
|  | Tone D (294 Hz) | 2 |
|  | Tone E (330 Hz) | 2 |
| **Mixture** | **Square + Auditory** |  |
|  | Square + Vowel /a/ | 2 |
|  | Square + Vowel /o/ | 2 |
|  | Square + Vowel /i/ | 2 |
|  | Square + Tone C (262 Hz) | 2 |
|  | Square + Tone D (294 Hz) | 2 |
|  | Square + Tone E (330 Hz) | 2 |
|  | **Male Face + Auditory** |  |
|  | Male Face + Vowel /a/ | 1 |
|  | Male Face + Vowel /o/ | 1 |
|  | Male Face + Vowel /i/ | 1 |
|  | Male Face + Tone C (262 Hz) | 1 |
|  | Male Face + Tone D (294 Hz) | 1 |
|  | Male Face + Tone E (330 Hz) | 1 |
|  | **Female Face + Auditory** |  |
|  | Female Face + Vowel /a/ | 1 |
|  | Female Face + Vowel /o/ | 1 |
|  | Female Face + Vowel /i/ | 1 |
|  | Female Face + Tone C (262 Hz) | 1 |
|  | Female Face + Tone D (294 Hz) | 1 |
|  | Female Face + Tone E (330 Hz) | 1 |

**Fig. 3:**
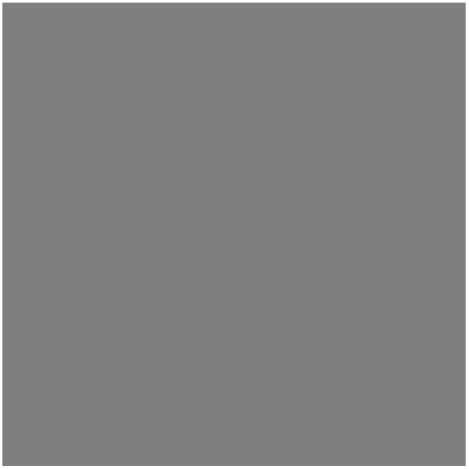
Example of Gray Square. Visual stimuli employed in the experiment, including a grayscale square.

### Data Preprocessing Pipeline

Preprocessing of the EEG data was carried out using EEGLAB v2022 ^31^ to ensure the integrity of the signal and the mitigation of artifacts. A causal FIR filter was used for bandpass filtering between 1 and 50 Hz, with a filter order of 500 and a buffer size of 30 seconds, effectively attenuating low-frequency drifts and high-frequency noise while preserving relevant neural oscillations. The data were subsequently resampled at 250 Hz to optimize computational efficiency while maintaining the fidelity of the recorded neural activity.

To address non-neural artifacts, artifact subspace reconstruction (ASR) ^32^ was applied with a threshold parameter as suggested in ^33^, which adaptively detects and reconstructs segments exhibiting excessive deviation from the statistical distribution of clean EEG data. The choice of the threshold parameter in ASR is essential for balancing between high-amplitude artifact removal and signal preservation, ensuring that transient high-amplitude artifacts (e.g. muscle activity, electrode displacement) are mitigated without excessively attenuating valid neural signals. Lower k values would result in overly aggressive rejection of artifacts, potentially removing informative neural activity, while higher values might allow significant artifacts to persist.

Following high-amplitude artifact correction, independent component analysis (ICA) ^34^ with linear interpolation was performed to decompose the multichannel EEG signals into statistically independent sources, facilitating the isolation of neural components from non-neural artifacts. Finally, ICLabel ^35^, a machine learning-based independent component classification tool, was utilized to automatically identify and remove artifacts related to ocular and muscular activities. Components classified as artifacts with a confidence probability greater than 0.8 were excluded from further analysis. This procedure effectively improved the SNR, ensuring that the retained components predominantly reflect neural activity.

### Data Records

The raw EEG recordings, stored in the BIDS-compliant format, have been publicly released on OpenNeuro (https://openneuro.org/datasets/ds005815/versions/1.0.1). The original data, initially in CNT format, were converted to BIDS using the MNE-Python and pybv packages.

### Technical Validation

To ensure the reliability and validity of the dataset, we conducted both behavioral and neurophysiological analyzes. Subjective vividness ratings were assessed to evaluate participants’ self-reported imagery experiences under different stimulus conditions. In parallel, neural responses were examined using event-related potentials (ERPs) to capture time-locked brain activity and power spectral density (PSD) analysis to characterize frequency-domain neural oscillations. These analyses provide complementary insights into how different sensory modalities influence mental imagery and cognitive processing.

### Vividness Ratings Analysis

Figure 4 presents the distribution of vividness ratings across different stimulus conditions. The results show that multimodal stimuli generally received higher and more consistent vividness ratings, while pure auditory stimuli exhibited greater variability. A three-way repeated measures ANOVA was conducted to analyze the effects of subject, stimulus condition, and session. Significant main effects were observed for subject (*F* = 239.54, *p <* .001), stimulus condition (*F* = 4.04, *p <* .001), and the session (*F* = 14.61, *p <* .001), indicating that the vividness varied between individuals, the stimulus conditions and the sessions. Interaction effects between the stimulus condition and the subject (*F* = 3.51, *p <* .001) and the subject and the session (*F* = 16.21, *p <* .001), while the interaction between stimulus condition and session (*F* = 0.67, *p* = .896)was not significant. A significant three-way interaction (*F* = 1.16, *p* = 0.009) was also observed.

**Fig. 4:**
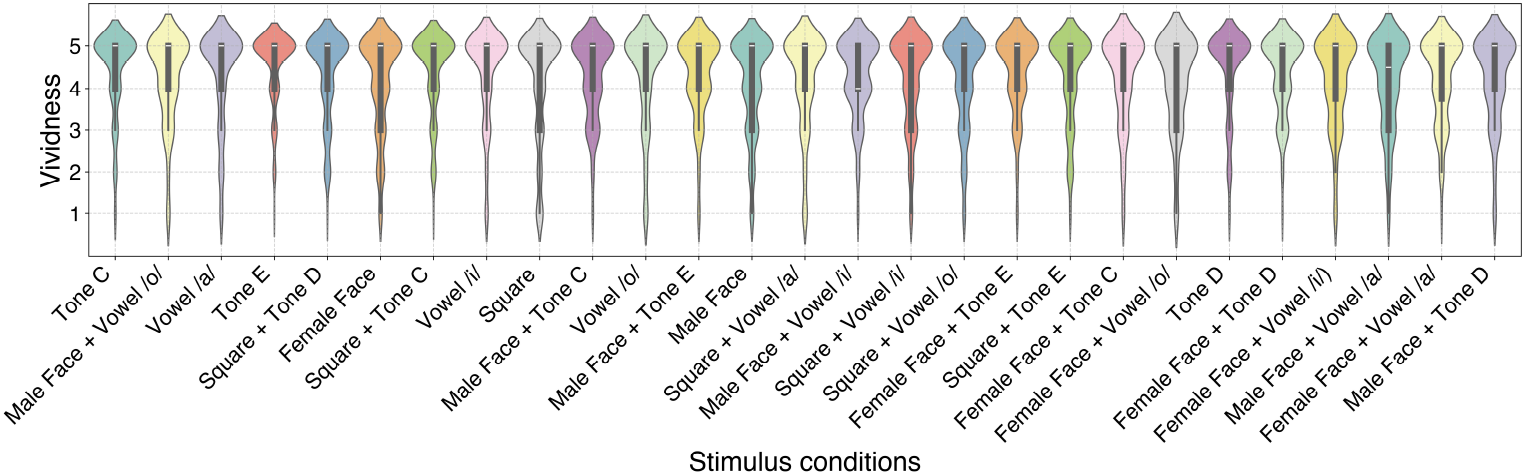
Distribution of Subjective Imagery Vividness Ratings Across Stimulus Conditions. Violin plot depicting the distribution of participant-rated vividness of mental imagery across different stimulus conditions. The horizontal axis categorizes auditory-only, visual-only, and multimodal stimuli, while the vertical axis indicates vividness ratings from 1 to 5.

### Event-Related Potentials (ERP) Analysis

Figure 5, 6, 7 show the ERP waveforms across stimulus conditions.

**Fig. 5:**
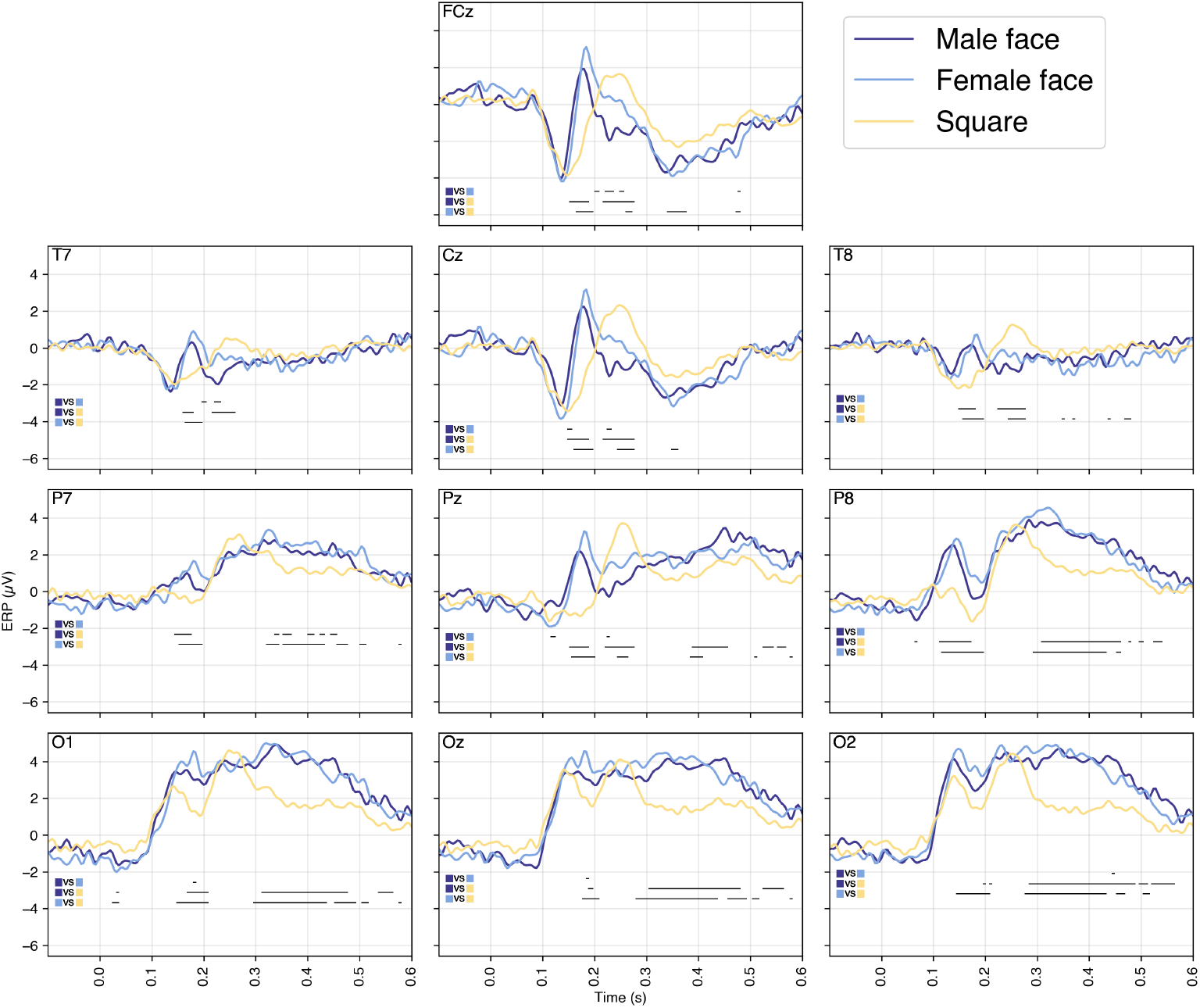
Event-Related Potentials (ERP) Waveforms for Visual Stimuli. ERP waveforms elicited by visual stimuli (male faces, female faces, and squares) at FCz, Cz, and Pz electrode sites. Facial stimuli evoke stronger neural responses compared to non-facial shapes, reflecting specialized neural processing of face perception.

**Fig. 6:**
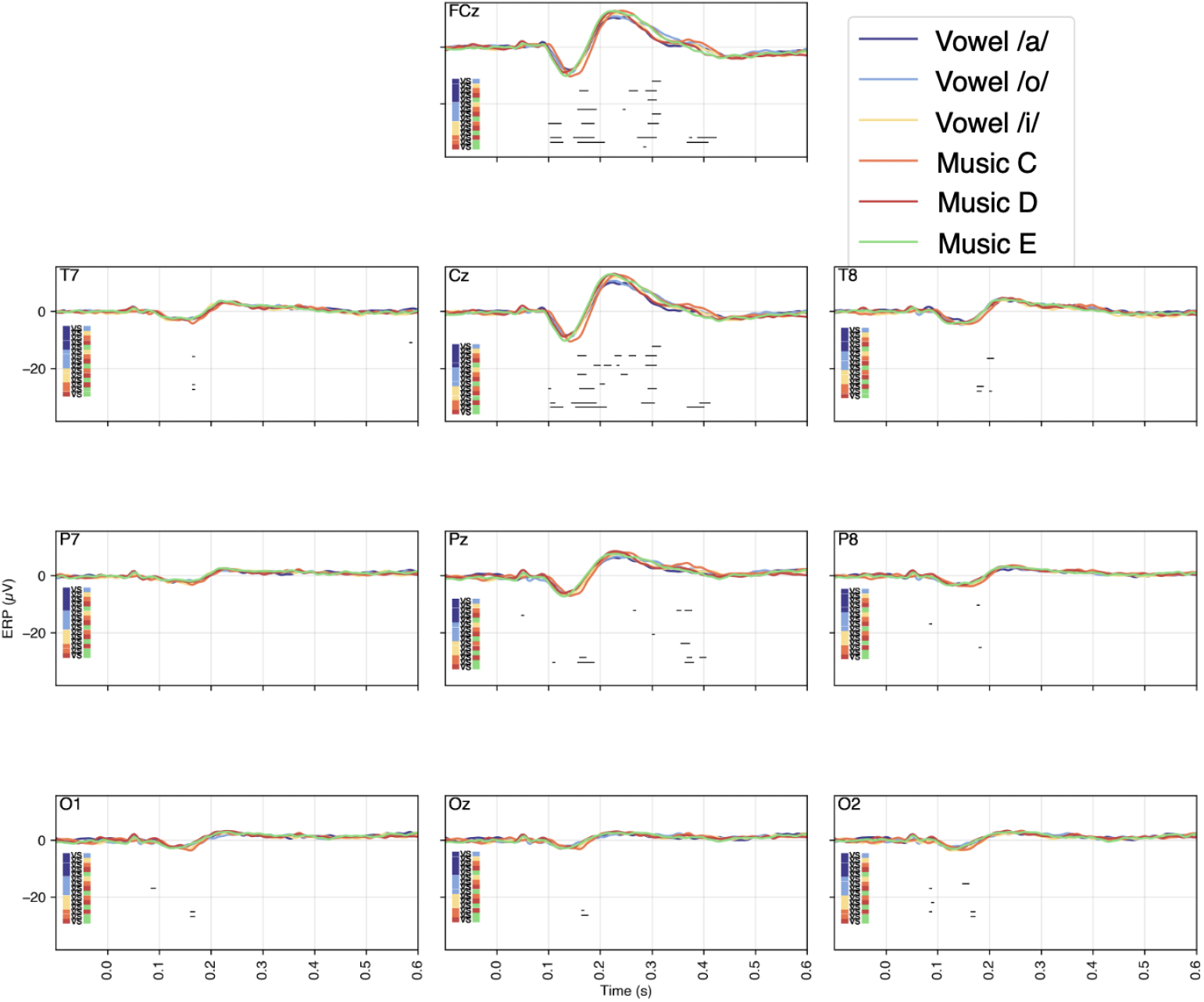
Event-Related Potentials (ERP) Waveforms for Auditory Stimuli. ERP waveforms comparing auditory stimuli responses (human vowels and musical tones) at different electrodes.

**Fig. 7:**
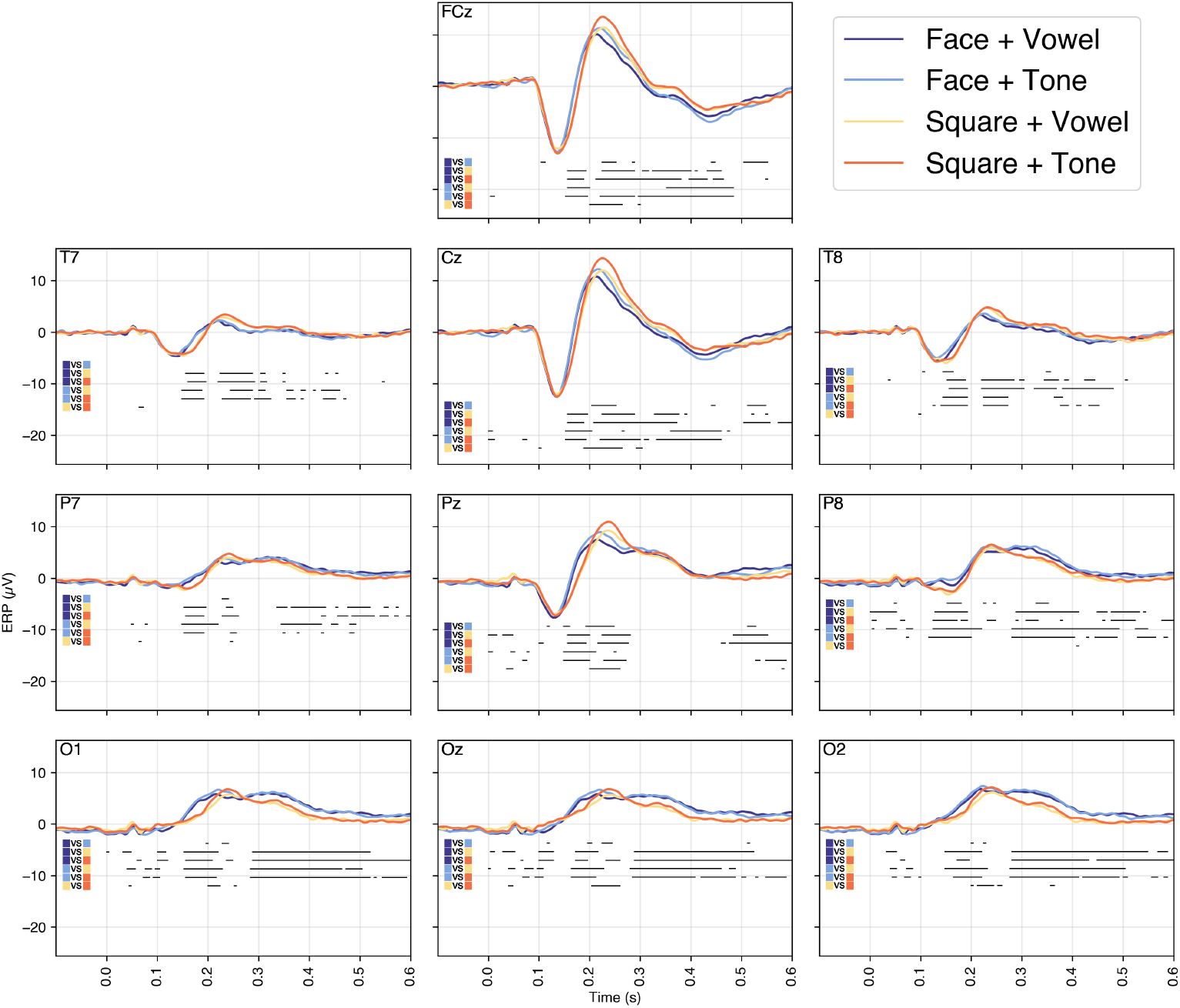
Event-Related Potentials (ERP) Waveforms for Multimodal (Visual + Auditory) Stimuli. ERP waveforms highlighting neural responses to combined visual and auditory stimuli.

The ERP of the visual stimuli, as illustrated in Figure 5, shows the waveform distributions of the visual stimuli, including male faces, female faces, and squares. The findings reveal that face stimuli evoke significantly higher amplitudes at the FCz, Cz, and Pz electrodes. These results highlight the role of neural networks involved in face processing, demonstrating greater sensitivity to facial stimuli compared to non-facial shapes ^36,37^.

The ERP of the auditory stimuli, depicted in Figure 6, compare neural responses to vocal and musical stimuli. The data indicate that vowels elicit higher amplitudes during early sensory responses (0.1–0.2 seconds), reflecting the brain’s rapid response to language-related sounds. In contrast, musical stimuli engage higher-level cognitive processes, such as emotional and melodic interpretation ^38–40^.

The ERP of the mixture stimuli, shown in Figure 7, illustrate the waveform characteristics of combined visual and auditory stimuli. The results demonstrate how sensory modalities are integrated, highlighting the brain’s ability to process and merge information from multiple sensory channels ^41–43^.

When comparing the differences in power spectral density (PSD) of brain waves between various types of imagined activities and the baseline condition (Fixation) (Figure 8), significant regional brain wave characteristics were observed in mixed visual, auditory and audiovisual imagery.

**Fig. 8:**
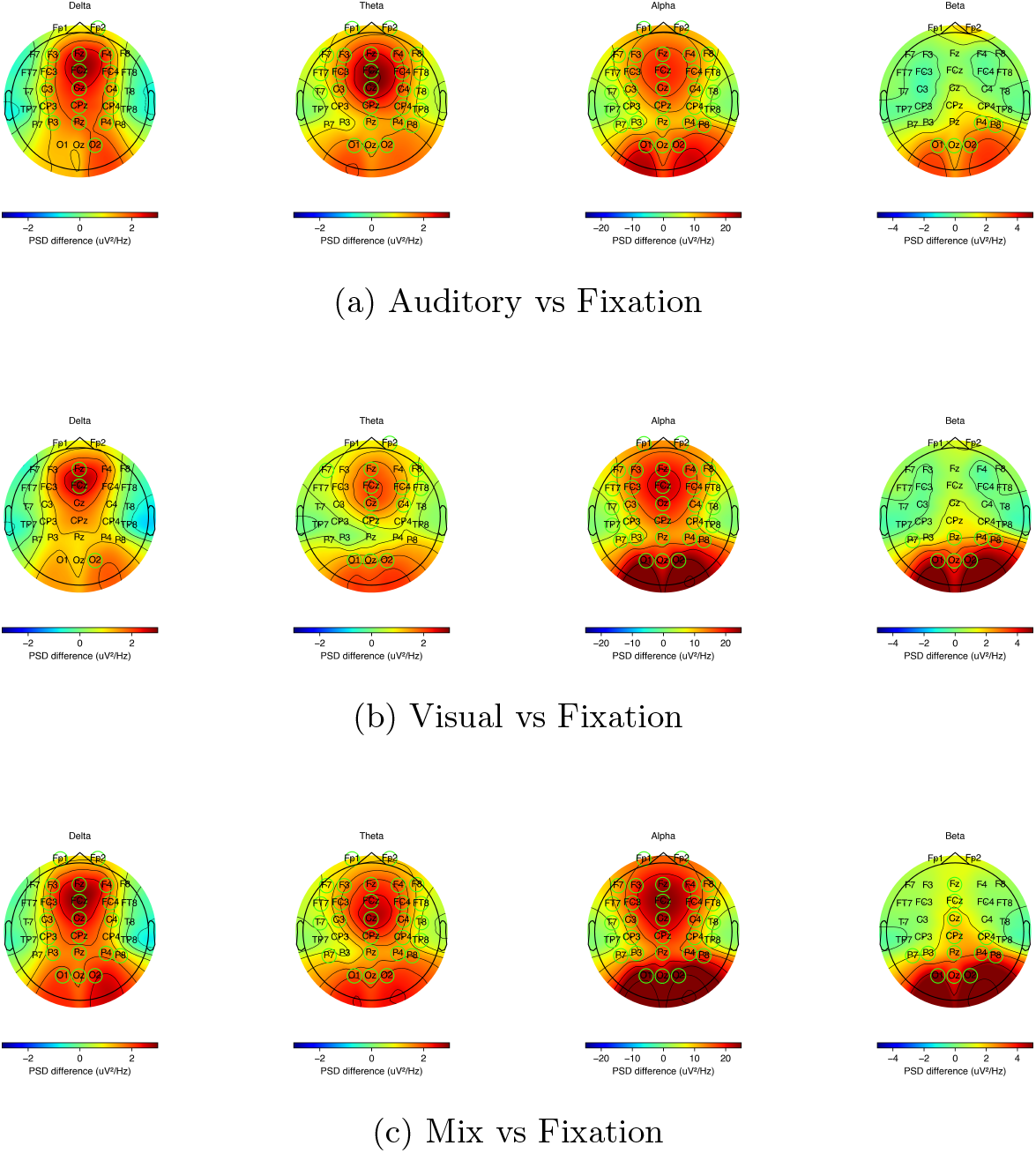
Power Spectral Density (PSD) Differences b etween I magery Tasks and Baseline Fixation. Comparative PSD analysis illustrating EEG spectral changes between imagery conditions (visual, auditory, multimodal) and baseline fixation. Visual imagery significantly enhances Alpha oscillations in the occipital lobe; auditory imagery primarily boosts Theta activity centrally; multimodal imagery broadly elevates neural oscillations across frequency bands.

Visual imagery (Figure 8a) primarily activated the enhancement of alpha wave responses in the occipital lobe, while auditory imagery (Figure 8b) showed a more pronounced enhancement of Theta waves in the central, frontal and parietal regions. Mixed audiovisual imagery (Figure 8c) showed significant changes in all frequency bands, with wave enhancements that significantly exceeded those of single-modality stimuli, further confirming that multimodal integration requires higher cognitive resources and integration of the brain network.

In Figure 9a, it can be seen that Delta and Theta waves show significant reductions in the frontal region, with the reduction in Theta waves being more prominent. In contrast, the Alpha and Beta waves exhibit significant enhancements in the occipital lobe (O1, O2, Oz).

**Fig. 9:**
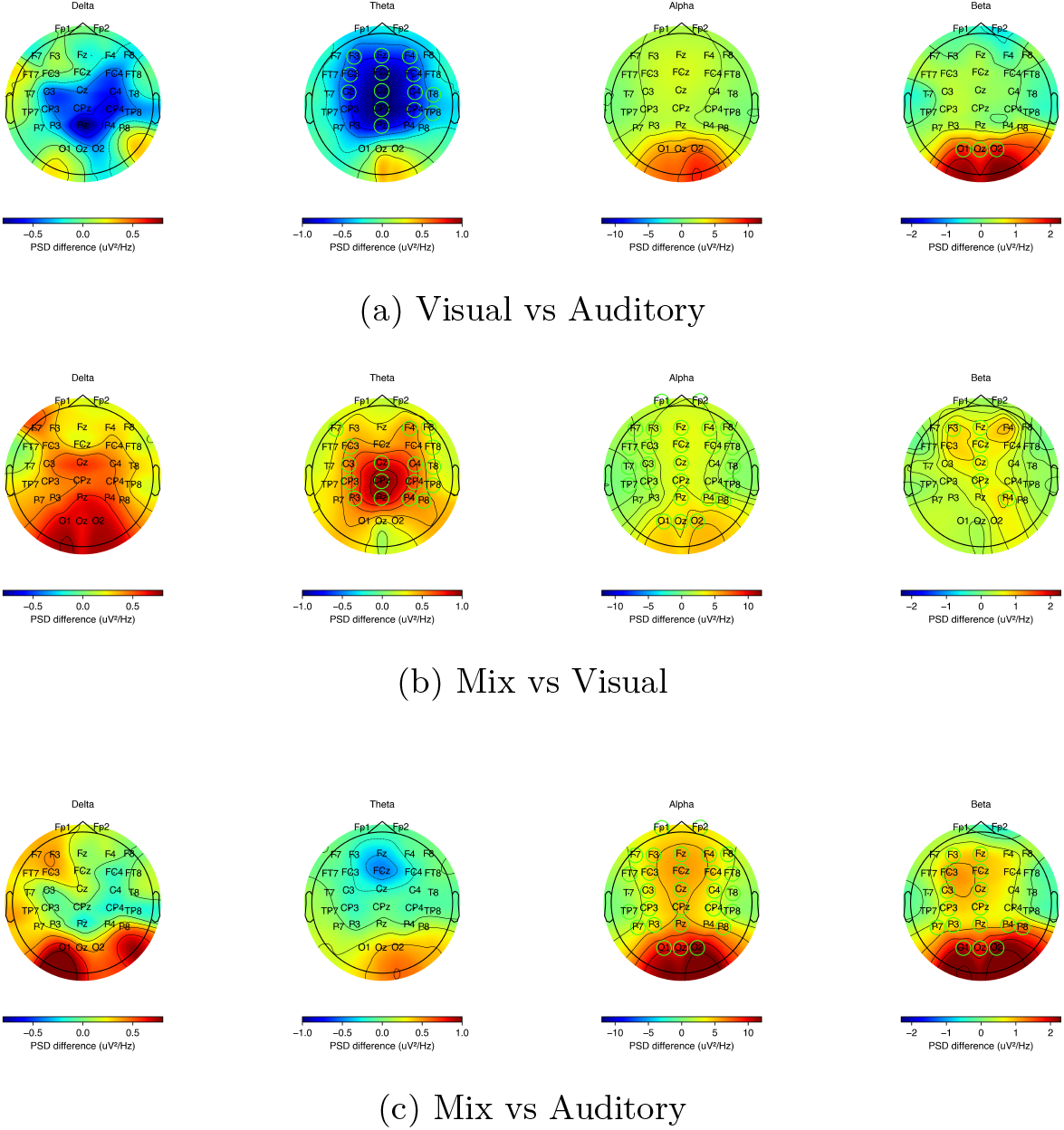
Power Spectral Density (PSD) Differences Among Various Imagery Conditions. Comparative PSD analyses highlighting neural spectral differences between (a) visual versus auditory imagery, (b) multimodal versus visual imagery, and (c) multimodal versus auditory imagery. Results underscore distinctive neural activation patterns across different brain regions and frequency bands, notably heightened Alpha and Beta activities in multimodal imagery.

In Figure 9b, the Delta waves show a slight enhancement in the occipital lobe, while the Theta waves demonstrate a significant improvement in the central (Cz) and parietal (Pz) regions. Alpha waves also show slight enhancement in the occipital lobe, while beta waves exhibit enhancement in the frontal and central regions, further confirming that mixed stimuli increase the demand for higher cognitive resources ^44–47^. In Figure 9c, the Delta waves show an improvement in the occipital lobe, with negligible differences in the central and frontal regions. Theta waves demonstrate significant enhancement in the central region, with minimal differences in the frontal region. Alpha waves show significant enhancement in the occipital and parietal regions, while beta waves exhibit significant enhancement in the frontal and central regions without any noticeable reductions, confirming the generally stronger Beta wave activity in multimodal stimuli ^48–50^.

These findings clearly validate the effectiveness of the data and demonstrate the significance of the YOTO data set in providing support for the EEG signal for both the resting state and the task phases, allowing the exploration of the random cognitive transition mechanisms of the brain.

## Code Availability

The source code used for all technical validations conducted in this experiment has been uploaded and is publicly available on GitHub (https://github.com/CECNL/YOTO_You_Only_Think_Once).

## Acknowledgements

This research was supported in part by the National Science and Technology Council (109-2222-E-009-006-MY3, 112-2321-B-A49-012 and 112-2222-E-A49-008-MY2); in part by the Healthy Longevity Global Grand Challenge Catalyst Award of National Academy of Medicine, USA, and Academia Sinica, Taiwan (AS-HLGC-113-06); in part by the National Health Research Institute, Taiwan (Grant NHRI-EX114-11418EC); and in part by the Higher Education Sprout Project of National Yang Ming Chiao Tung University and Ministry of Education.

## Author Contributions

Chun-Shu Wei: Conceptualization, Methodology, Investigation, Formal analysis, Visualization, Data Curation, Writing – Original Draft, Writing – Review & Editing, Supervision, Project administration. Yan-Han Chang: Formal analysis, Investigation, Visualization, Data Curation, Writing – Original Draft, Writing – Review & Editing. Hsi-An Chen: Formal analysis, Investigation, Visualization, Writing – Original Draft, Data Curation. Min-Jiun Tsai: Methodology, Investigation, Data Curation, Project administration. Chun-Lung Tseng: Investigation, Data Curation, Formal analysis. Ching-Huei Lo: Investigation, Data Curation. Kuan-Chih Huang: Methodology, Data Curation, Project administration.

